# An RNA regulates iron homeostasis and host mucus colonization in *Bacteroides thetaiotaomicron*

**DOI:** 10.1101/2025.09.08.672848

**Authors:** Gianluca Prezza, Ryan T. Fansler, Thomas Guest, Gohar Mädler, Hannah Schlauch, Wenhan Zhu, Alexander J. Westermann

## Abstract

Symbiotic bacteria in the human intestinal microbiota provide many pivotal functions to human health and occupy distinct biogeographic niches within the gut. Yet the molecular basis underlying niche-specific colonization remains poorly defined. To address this, we conducted a time-resolved dual RNA-seq experiment to simultaneously monitor the transcriptional co-adaptations of human commensal *Bacteroides thetaiotaomicron* and human gut epithelial cells in an anaerobe-epithelium co-culture system. Comparative transcriptomic analysis of mucus-associated versus supernatant *Bacteroides* populations unveiled small RNAs (sRNAs) that are differentially regulated between spatially segregated subpopulations. Among these, we identified IroR as a key sRNA that facilitates *B. thetaiotaomicron* adaptation to the mucus-rich, iron-limiting niche, partly by modulating expression of bacterial capsule genes. This work provides new insights into the spatiotemporal dynamics of gut colonization and underscores a previously underappreciated role for bacterial sRNAs in shaping mutualistic interactions between the human microbiota and the gut epithelium.

## INTRODUCTION

The gastrointestinal tract is home to trillions of microbes, which together form a complex community that provides many pivotal functions to human health [1]. These commensal bacteria are distributed along the entire gut, with density and diversity rising along its longitudinal axis [2]. Within the large intestine, a combination of physicochemical gradients and mucous layer composition creates distinct microenvironments between the epithelial surface and the lumen [3, 4]. Many bacterial species, such as *Bacteroides thetaiotaomicron*, occupy multiple intestinal niches, including the mucous layer, food particles, or the intestinal lumen [5, 6]. To thrive in these varied habitats, *Bacteroides* spp. have evolved intricate mechanisms, including an extensive repertoire of polysaccharide utilization loci (PULs), enabling them to efficiently forage complex host-and diet-derived glycans and competitively colonize these gut niches [7].

The specific intestinal niches occupied by commensal *Bacteroides* can have profound consequences on the human host’s health. In healthy individuals, these commensals typically ferment dietary fibers and can promote intestinal homeostasis by educating the host immune system and providing colonization resistance against enteric pathogens [8, 9]. However, when deprived of dietary fibers, *Bacteroides* will instead consume host-secreted glycoproteins and erode the intestinal mucous layer, compromising the defense against enteric pathogen infection [10]. Additionally, mucus utilization by *B. thetaiotaomicron* has been shown to increase intestinal inflammation and exacerbate immunological disorders, such as graft-versus-host disease [11]. Thus, understanding the regulatory mechanisms that govern *B. thetaiotaomicron*’s adaptation to the mucous layer niche is essential for advancing human health.

The mucous layer presents a distinct nutritional landscape, with host-derived, rather than dietary, glycans predominating as the primary carbon source [12]. Additionally, bioavailable iron levels are markedly lower in the mucous layer compared to the lumen, likely due to increased levels of oxygen, which oxidize soluble iron (II) to insoluble iron (III) [13, 14]. Successful colonization of the mucous layer, therefore, requires bacteria to tightly regulate acquisition programs to compete for these essential nutrients [13–15]. While transcriptional and post-transcriptional regulation involved in the lifestyle of *B. thetaiotaomicron* in the human gut have recently begun to be elucidated [16, 17], significant knowledge gaps remain. Several studies have investigated *Bacteroides* transcriptomic activity in mouse models of colonization. However, the inherent complexity of animal experiments often limits these studies to characterizing only a single time point during colonization [18–22] or to collecting bacterial samples only from the luminal contents [23]. These limitations hinder our understanding of the spatiotemporal dynamics of colonization and bacterial behavior near the host mucosa.

Despite major advances in understanding bacterial adaptation in the gut, post-transcriptional regulation by small RNAs (sRNAs) remains largely unexplored in commensal mutualists. sRNAs are established regulators of stress responses and host adaptation in pathogenic bacteria [24–27], but their roles in shaping the mutualistic behaviors of dominant commensals like *B. thetaiotaomicron* are only beginning to be appreciated. We recently identified hundreds of sRNAs in *B. thetaiotaomicron* [28, 29], some of which may play a previously unappreciated role during colonization of the mucous layer [16]. For instance, the FopS (for family of paralogous sRNAs) cluster, in concert with RNA-binding protein RbpB, has been implicated in polysaccharide utilization and mucus consumption in the mouse gut [30]. Additionally, sRNAs are well-established regulators of iron homeostasis across diverse bacterial species [31–35], making them strong candidates for modulating adaptation to the iron-limited mucous layer.

In addition to the *Bacteroides* adaptation to the epithelial niche, the reciprocal host epithelial response to *Bacteroides* colonization also remains incompletely understood. The polysaccharide A (PSA) of *B. fragilis* has long been appreciated for its immunomodulatory induction of IL-10 and regulatory CD4^+^ and CD8^+^ T cells [36, 37]. Recent studies have further demonstrated that *Bacteroides*-derived metabolites, including propionate [38], sphingolipids [39], and secondary bile acids [40], promote goblet cell differentiation, increase expression of genes involved in mucus production, and suppress pro-inflammatory signaling. Despite these advances, a time-resolved study to capture how the dynamic host epithelium response directs *Bacteroides* colonization has yet to be presented. Such data could reveal how host responses to *Bacteroides* evolve over time, offering key insights into the mechanisms that maintain gut homeostasis.

Here, we conducted a time-resolved dual RNA-sequencing experiment to profile *B. thetaiotaomicron* as it colonizes a mucus-producing *in vitro* model of the human gut epithelium. This approach allowed us to simultaneously track the gene expression dynamics of host and bacteria, revealing their coordinated responses during colonization. Among other observations, we identified *B. thetaiotaomicron* sRNAs involved in these bacterial interactions with the mucous layer. We validate one such sRNA, the <u>Iro</u>n-responsive <u>R</u>NA (IroR, locus tag BTnc022), as a factor regulating colonization of the mucous layer in a context-dependent manner. Functional characterization of IroR revealed its role in adapting *B. thetaiotaomicron* to iron limitation and in promoting the utilization of host-derived polysaccharides, in part by regulating capsule gene expression. These regulatory effects manifest in measurable phenotypic outcomes in the murine gut.

## RESULTS

### Establishing a hypoxic, mucus-producing epithelial host model of B. thetaiotaomicron colonization

Mucous layer colonization by resident commensals profoundly impacts host physiology, yet the host and bacterial responses remain poorly understood. To address this, we developed a Transwell-based cell culture system that models the commensal-epithelial interactions *in vitro*. HT29-MTX-E12 (from here on, HT29) cells, an immortalized human cell line differentiated into a goblet-like phenotype [41], were seeded on the apical side of the Transwell. These cells were then cultured under normoxic conditions to facilitate cell layer tightening, cell polarization, and mucus secretion, as shown previously [42–44]. After 28 days of maturation, the HT29 cells successfully divided the Transwell into a mucus-secreting apical compartment and a basolateral compartment (**Fig. 1A**). Positive staining for gel-forming Mucin-2 (MUC2) (**Fig. 1B, left**) confirmed the presence of mucus-secreting goblet-like cells, and the expression of tight junction markers, such as Zonula Occludens-1 (ZO1), indicating that the cellular monolayer had achieved classic tight barrier characteristics. Altogether, this shows that our Transwell system models key features of the gut epithelial layer *in vitro*.

**Figure 1:**
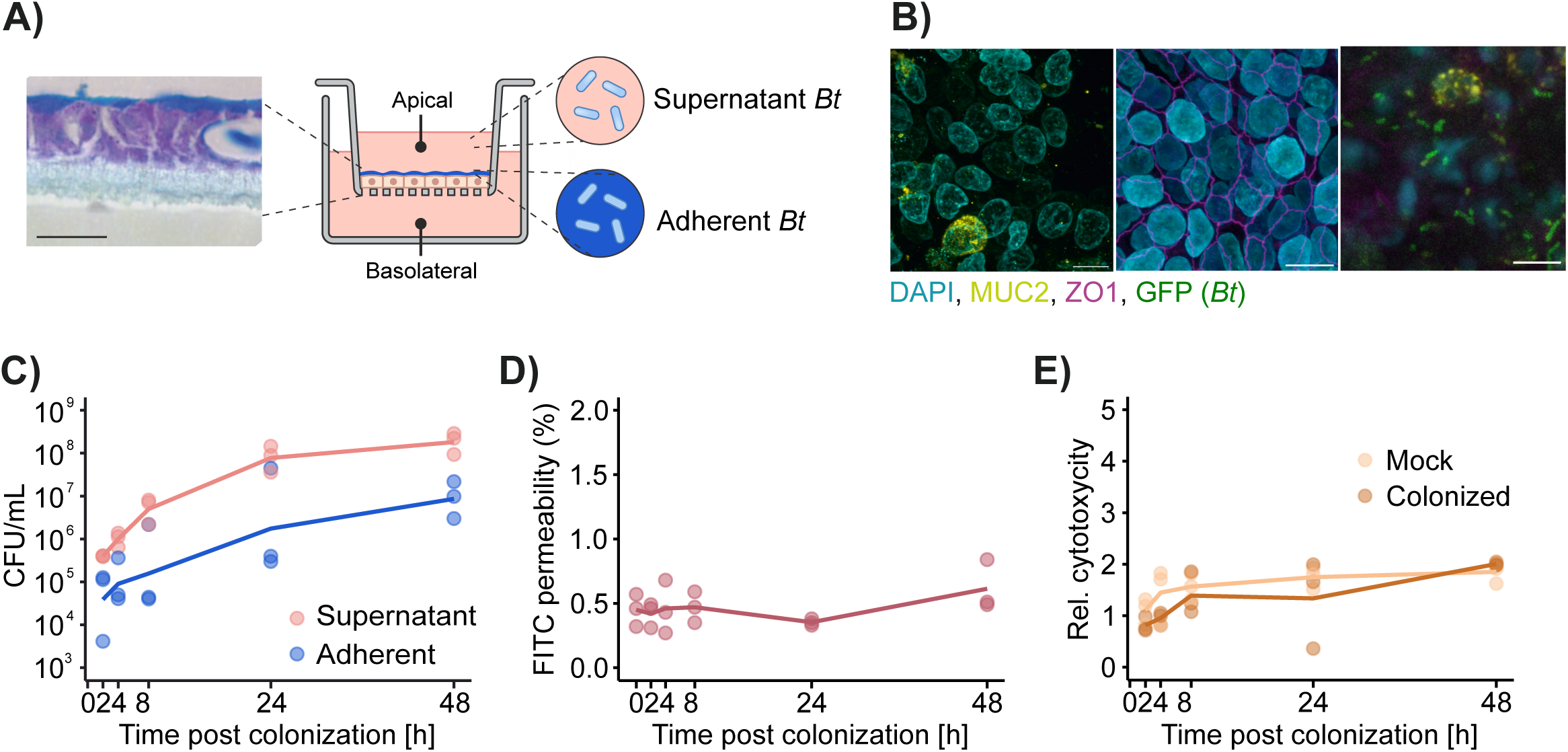
Colonization of a cell-culture model of the gut epithelium. A) Human cells are seeded on the semipermeable membrane of the Transwell system and mature for 28 days, until the cell layer is tight and a mucus layer is formed. Left: hematoxylin-eosin and alcian blue staining of a cross-section of a mature Transwell model. Mucus is stained in light blue, cells are purple, and the porous polyester membrane is transparent. Subpopulations of *B. thetaiotaomicron (Bt)* establish in the supernatant and adherent to the mucous layer. Scale bar: 25 μm. B) Immunofluorescence staining of secreted mucin MUC2, the tight junction marker ZO1 and GFP-expressing wild-type *B. thetaiotaomicron*. Shown are representative top-view images of the mature Transwell model. Scale bars: 10 µm. C) Colonization of the model by *B. thetaiotaomicron*. Plotted are the CFU per ml in the supernatant and adherent fractions of the Transwell. A line represents the mean value (n=3). D) Integrity of the epithelial barrier over the course of colonization. Data show the percentage of fluorescence detected in the basolateral compartment after addition of FITC-glucose to the apical side. A line represents the mean value (n=3). E) Host cytotoxicity assay. Activity of the cytosolic lactate dehydrogenase is measured in the apical medium and normalized to the timepoint 0. A line represents the mean value (n=3).

We next incubated the model under 1% O_2_ atmospheric concentration to better simulate conditions in the colon [45]. *B. thetaiotaomicron* cells were added to the host medium in the apical compartment at a multiplicity of colonization (MOC) of 10 bacterial cells per host cell. Colonization was successfully sustained over the span of 48 h; bacterial colony-forming unit (CFU) counts from both within the host mucous layer (“adherent” subpopulation) and in the supernatant medium (“supernatant”) increased over time (**Fig. 1C**), similar to the distinct intestinal niches in the mouse colon [21]. The epithelial barrier integrity (**Fig. 1D**) and host cell viability (**Fig. 1E**) were not affected by *B. thetaiotaomicron* colonization. Together, we have established an amenable cell culture model suitable for studying the regulatory processes that underlie host colonization by this anaerobic gut commensal.

### Dual RNA-seq of B. thetaiotaomicron-colonized gut epithelial cells

Having established an *in vitro* system to model the gut epithelial niche, we next sought to elucidate the molecular activities that underpin *Bacteroides* colonization of the host mucous layer. The colonized Transwell model allowed us to simultaneously profile both host and bacterial transcriptomes via dual RNA-seq to uncover regulatory activities at the interface of the epithelial layer [46, 47]. The dual RNA-seq method has been widely used to investigate infection by pathogenic bacteria both in cell culture models (e.g., [48–50]) and *in vivo* (e.g. [51–54]). However, to date, dual RNA-seq has not been used to study host colonization by a commensal bacterium.

*B. thetaiotaomicron* does not directly adhere to host cells, but dwells within the mucous layer [5]. As a result, commonly used methods to enrich bacteria in direct contact with host cells, such as fluorescence-activated cell sorting [48, 52, 55], are not amenable to isolating mucus-associated *B. thetaiotaomicron*. We therefore carefully collected the supernatant without damaging the mucosal lining and extracted total RNA separately from both adherent and supernatant fractions. We estimated by qRT-PCR that bacterial RNA collected from the adherent fraction of a colonized Transwell would account for ∼0.3% and ∼3% of the total RNA at 2 and 48 h post-colonization (p.c.), respectively (**Suppl. Fig. S1A**). As inferred from previous experience [56], it would thus be possible to generate sequencing libraries with coverage of the *B. thetaiotaomicron* transcriptome sufficient for differential gene expression analysis, despite the relative abundance of host RNA in the adherent fractions.

We thus designed a dual time-course RNA-seq experiment, including three biological replicates per condition. Before colonization, total RNA was extracted from the input bacterial cultures and uncolonized host cells (time point 0). Then, RNA was extracted from adherent and supernatant fractions of the Transwell model at 2, 4, 8, 24, and 48 h p.c., in addition to mock-treated host cells. After depletion of host and bacterial ribosomal RNAs (rRNAs), we generated and sequenced cDNA libraries (**Suppl. Fig. S1B**). The fraction of total reads mapping to the *B. thetaiotaomicron* genome in the adherent samples steadily increased over time, from ∼0.1% at 2 h p.c. to ∼20% at 48 h p.c. (**Suppl. Fig. S1C**, Table 2). This increase in mapped reads accords with the bacterial CFU counts over the same time scale (**Fig. 1C**). As expected, reads obtained from the supernatant fraction were mostly of bacterial origin (Fig. S1C), with a small proportion of human-derived reads likely from dead, lysed host cells. Host-derived reads were efficiently depleted of rRNA and were dominated by mRNAs and long noncoding RNAs (**Suppl. Fig. S1D**). Although rRNA depletion for *B. thetaiotaomicron* was less efficient, most bacterial reads aligned to informative genes, with ∼65% to mRNAs and ∼6% to sRNAs (**Suppl. Fig. S1E**). In summary, the dual RNA-seq dataset is of high quality and well-suited for identifying differentially expressed genes in both host and microbe during their interaction.

### The host mounts a tolerant response to colonization by B. thetaiotaomicron

HT29 cells basally express various classical pathogen recognition receptors (PRRs) [57–60]. Nevertheless, there were no differentially expressed genes in time-matched *B. thetaiotaomicron*- and mock-colonized host cells at early time points (2, 4, or 8 h p.c.) (**Suppl. Fig. S2**). This is in contrast to epithelial cells infected by the enteric pathogen *Salmonella* Typhimurium, in which host responses can be readily detected much earlier at 2 h post-infection [61]. To get a comprehensive overview of transcriptional changes in response to *B. thetaiotaomicron* at the later time points, we organized the gene ontology (GO) terms and Kyoto Encyclopedia of Genes and Genomes (KEGG) pathways of differentially regulated genes in two enrichment maps [62, 63], where gene sets sharing common genes are binned in clusters (**Fig. 2**). At 24 h p.c., the most prevalent transcriptional signature in colonized HT29 cells was an upregulation of endoplasmic reticulum (ER) stress response, protein folding, and Golgi apparatus gene sets (**Fig. 2A**). This is consistent with previous studies, which have shown that *B. thetaiotaomicron* stimulates the secretion of mucus from goblet cells [64–66]. The ER stress response at 24 h p.c. could be triggered by this increase in mucus production. However, the *Muc2* mRNA, encoding for a major protein component of colonic mucus [67], was not differentially expressed at any time (Fig. S2), and we speculate that *B. thetaiotaomicron* may induce mucus secretion independently of transcriptional control of *Muc2*.

**Figure 2:**
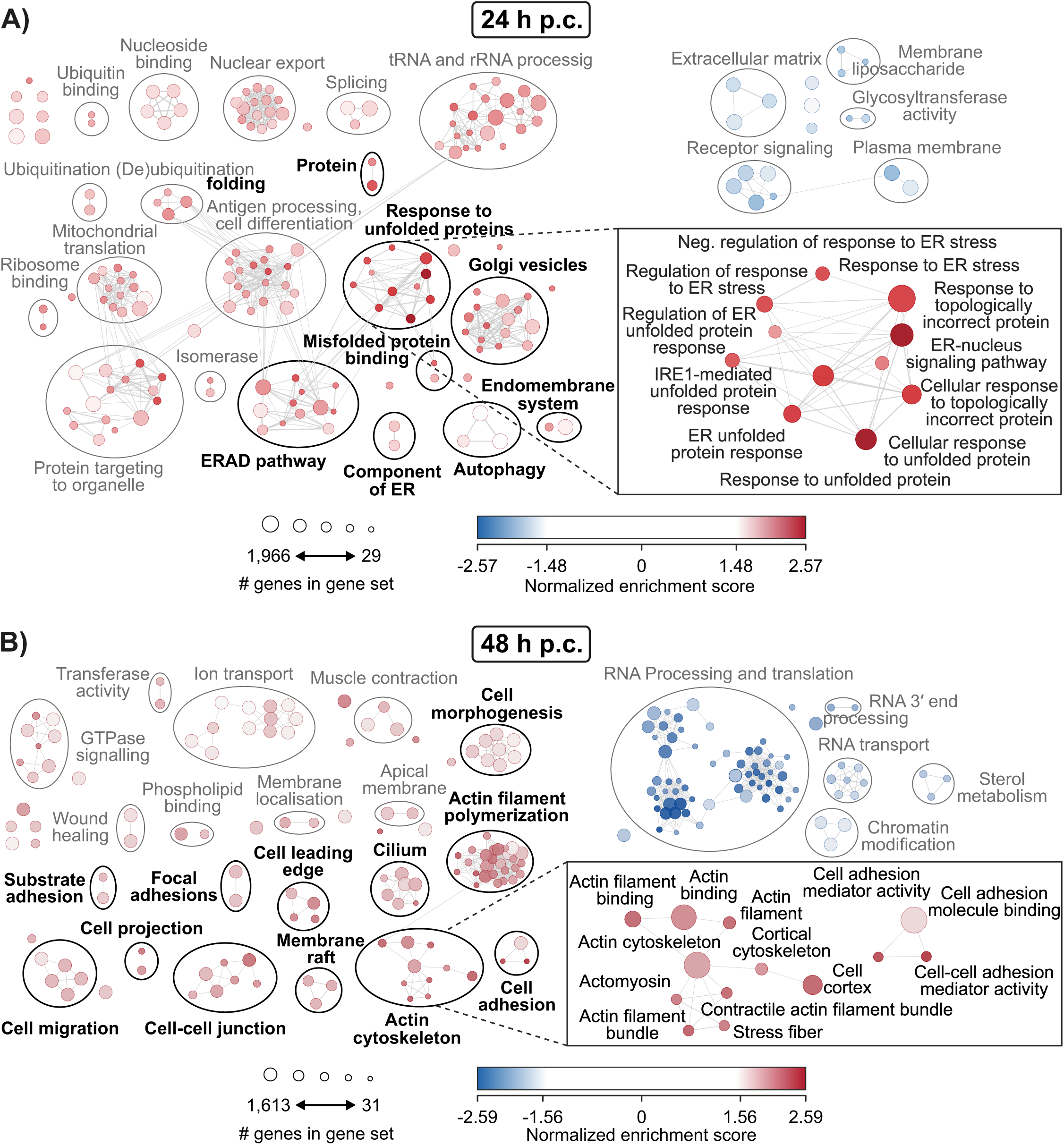
The host response to colonization. A) Enrichment map of significantly (FDR < 10^-3^) (de)enriched host gene sets 24 hours post-colonization. Each gene set is represented as a circle of size proportional to the number of genes it includes. The fill color of the circles is determined by the normalized enrichment score of each gene set. A positive score indicates upregulation of the set in colonized cells. Gene sets sharing genes are connected by lines, the width of which is proportional to the number of overlapping genes. Sets are clustered with Autoannotate [121] and manually named. A box in the lower right corner shows some of the pathways involved in ER stress. B) Enrichment map of significantly (FDR < 10^-5^) (de)enriched host gene sets 48 hours post-colonization. A box in the lower right corner shows some of the pathways involved in cytoskeletal rearrangement. See panel A for a detailed description of the plot elements.

At 48 h p.c., colonized host cells upregulated various pathways involved in cytoskeletal rearrangement, cell adhesion, and junctions (**Fig. 2B**). The let-7 family of microRNAs was also downregulation in colonized models at this time point, similar to the response to infection by *S*. Typhimurium [61], albeit much later (**Suppl. Fig. S2**). Let-7 represses the immunomodulatory cytokines IL-6 and IL-10, and is downregulated upon extracellular lipopolysaccharide (LPS) sensing by Toll-like receptor-4 (TLR4) as part of the innate immune response [61]. Altogether, our Transwell model provides insights into the nature and dynamics of the epithelial response to colonization by commensal *B. thetaiotaomicron*, revealing clear distinctions from the response to pathogens. The relatively delayed and muted host immune response to commensal *B. thetaiotaomicron* compared to pathogens might be explained by cell surface structures that help *Bacteroides* evade detection by host PRRs, including different capsules (see below), and provides a unique opportunity to understand the commensal adaptation to host niches. In line with this reasoning, we therefore focused our efforts on elucidating bacterial regulatory features at the mucous layer.

### B. thetaiotaomicron adherent to the epithelium are metabolically active

To analyze the bacterial transcriptomic data, we identified pathways with the strongest transcriptional signatures, many of which are associated with bacterial features relevant during host colonization (**Fig. 3A**). We focused on those GO terms and KEGG pathways that are highly (|normalized enrichment score| > 2) and significantly (FDR < 0.05) (de)enriched in at least one time point (**Fig. 3B-i**). We observed upregulation of numerous pathways involved in carbon, energy, short-chain fatty acid (SCFA), and amino acid metabolism in adherent *B. thetaiotaomicron*, indicating that these bacteria are metabolically active. Of note, most of these processes are dependent on iron, a requirement that becomes increasingly challenging as microbes enter the mucous layer, where iron availability is limited [14]. We checked the expression levels of a list of genes known to be differentially expressed by *B. thetaiotaomicron* during growth in iron-deplete conditions [68, 69] (**Fig. 3C-i**). Indeed, significant up-regulation of xenosiderophore-encoding *xusB* and down-regulation of ferredoxin gene *BT_2414* in the adherent subpopulation are in accordance with this mucous-associated niche being depleted of iron, suggesting its successful colonization requires commensals to adapt to an iron-restricted environment.

**Figure 3:**
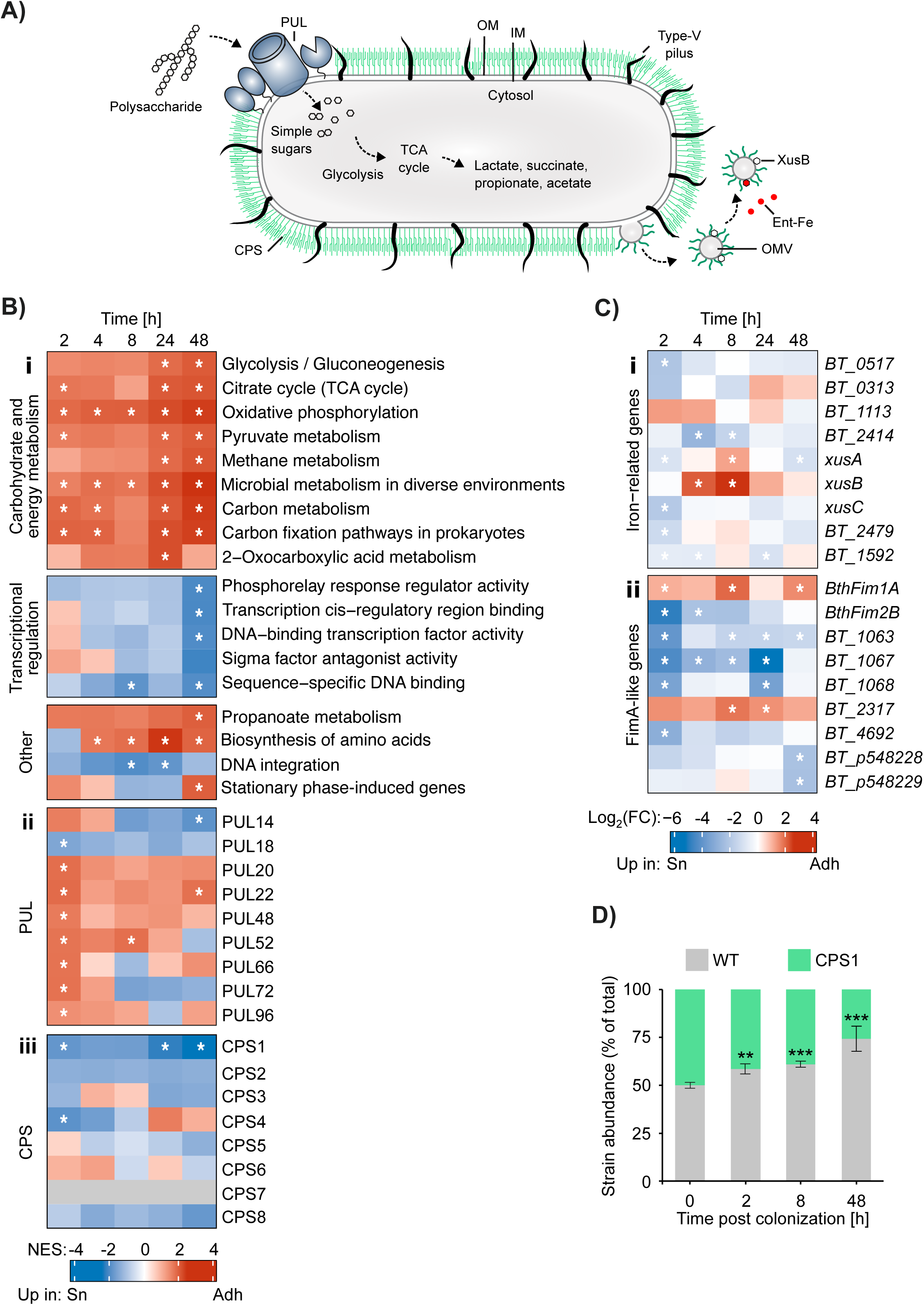
*B. thetaiotaomicron* transcriptional dynamics during interaction with the host. A) Schematic showing *B. thetaiotaomicron* molecular pathways relevant to host colonization. The polysaccharide utilization loci (PUL) allow acquisition and utilization of various polysaccharides as carbon sources. Capsular polysaccharides (CPS) and Type-V pili mediate host-microbe interactions. The XusABC iron acquisition system, present on outer membrane vesicles (OMV), sequester iron-laden xenosiderophores (such as Enterobactin). B) Dynamics of differentially regulated (i) GO Terms, KEGG pathways, (ii) PUL, and (iii) CPS gene sets between adherent and supernatant bacteria during colonization. NES: normalized enrichment score. An asterisk indicates FDR < 0.05. C) Upper panel (i) shows expression of selected genes known to be differentially expressed in iron-limited conditions, in adherent versus supernatant bacteria. Lower panel (ii) shows expression of type V pilus structural or assembly genes in adherent versus supernatant bacteria. An asterisk indicates FDR < 0.05. D) Competitive colonization experiment with a 1:1 mixture of WT (Δ*tdk*) and a strain expressing only CPS1. The abundance of each is determined by qRT-PCR amplification of unique genomic regions. The time point 0 represents the input strain mix.

Genes harboring a promoter associated with oxidative stress and stationary phase-induced genes (identified by the presence of the characteristic ‘promoter motif 2’ [29]) were enriched in the adherent subpopulation at 48 h (**Fig. 3B-i**). The expression of several PUL genes was generally higher in adherent than supernatant *B. thetaiotaomicron* (**Fig. 3B-ii**). Reassuringly, most of the identified differentially regulated PULs have previously been linked to the consumption of mucus-derived glycans (PULs 14, 18, 20, 22, 72 and 96) [70–72] or components of the extracellular matrix (PUL96) [73]. Relative expression of two such PULs (14 and 72) shifted over time, initially elevated at 2 and 4 h p.c. in the adherent bacteria, but later enriched in supernatant bacteria by 24 and 48 h p.c., possibly reflecting the continuous diffusion of cleaved mucin *O*-glycans into the supernatant. We also observed upregulation of starch (PUL66 = Sus) and dextran (PUL48) degradation PULs in adherent bacteria [68, 69]. Since starch and dextran are not present in the cell medium used, we hypothesize that the degradation of these dietary carbohydrates is transcriptionally coregulated with host glycan utilization, allowing *B. thetaiotaomicron* to be primed to consume other, preferred carbon sources should they become available.

In contrast to the adherent population, *B. thetaiotaomicron* in the supernatant upregulated many transcriptional regulators comprising classical and hybrid two-component systems and AraC-like transcription factors, especially at 48 h p.c. (Table 3). A large fraction of these regulators, including those belonging to annotated PULs, are upregulated in response to carbon starvation [28] (Table 3). We hypothesize that the supernatant population experiences carbon limitation by 48 h p.c. as glucose in the DMEM medium is depleted. In support of this, the activity of Cur, a transcriptional master regulator induced by carbon limitation [76, 77], is elevated in supernatant *B. thetaiotaomicron* compared to adherent bacteria at 48 h p.c. (**Suppl. Fig. S3A**).

### Selective expression of adhesion factors and capsule loci in mucus-associated bacteria

*Bacteroides* spp. express type-V pili, a distinct class of more than 30 adhesins that they share with other members of the *Bacteroidia* [78]. Nine of these *fimA*-like pili genes were differentially expressed between adherent and supernatant bacteria (**Fig. 3C-ii**). While the majority were more highly expressed in supernatant bacteria, *BthFim1A* and *BT_2317* were consistently upregulated in the adherent fraction, suggesting these could play a role in localization to the mucous layer.

The polysaccharide capsule of *B. thetaiotaomicron* is synthesized from one of eight CPS genomic loci, allowing these bacteria to alter their surface structure [79]. The capsule influences phage sensitivity [80], host immune evasion and modulation [81, 82], biofilm formation [83], and nutrient acquisition [84]. During early colonization, we observed downregulation of CPS4 in adherent bacteria, along with sustained repression of the CPS1 locus throughout the time course (**Fig. 3B-iii**). At 48 h p.c., CPS1 comprised a large portion of the most strongly downregulated genes in adherent compared to non-adherent bacteria (**Suppl. Fig. S3B**), indicating that its repression may be part of a niche adaptation strategy. Consistent with this notion, wild-type *B. thetaiotaomicron* outcompeted a strain in which expression of CPS1 is constitutively active (locked) [81] during competitive colonization of our Transwell model (**Fig. 3D**). These data complement previous findings [85] and inform about the cell-surface compositions that favor or hinder *B. thetaiotaomicron* adherence to mucus.

### Expression of B. thetaiotaomicron sRNAs during host colonization

Bacterial sRNAs are well established regulators of virulence and host interactions in pathogenic bacteria [26, 27], yet their roles in commensal species remain largely unexplored [86]. To investigate the role of *B. thetaiotaomicron* sRNAs during host colonization, we screened for sRNAs that were differentially expressed in adherent and supernatant bacteria. Among the 36 sRNAs that were significantly differentially expressed between the two populations in at least one time point (∼10% of the annotated sRNAs [28]), the uncharacterized candidate sRNAs BTnc022 (IroR = <u>Iro</u>n-responsive <u>R</u>NA, for reasons to follow), BTnc060, and BTnc182 were consistently upregulated in adherent bacteria (**Fig. 4A**). IroR and BTnc182 were among the most expressed genes in this population (**Suppl. Fig. S3C**). When comparing their levels to those in the input culture at time point 0 h, IroR and BTnc182 were upregulated up to ∼100-fold in adherent cells and only up to 30-fold in the supernatant population (**Fig. 4B**).

**Figure 4:**
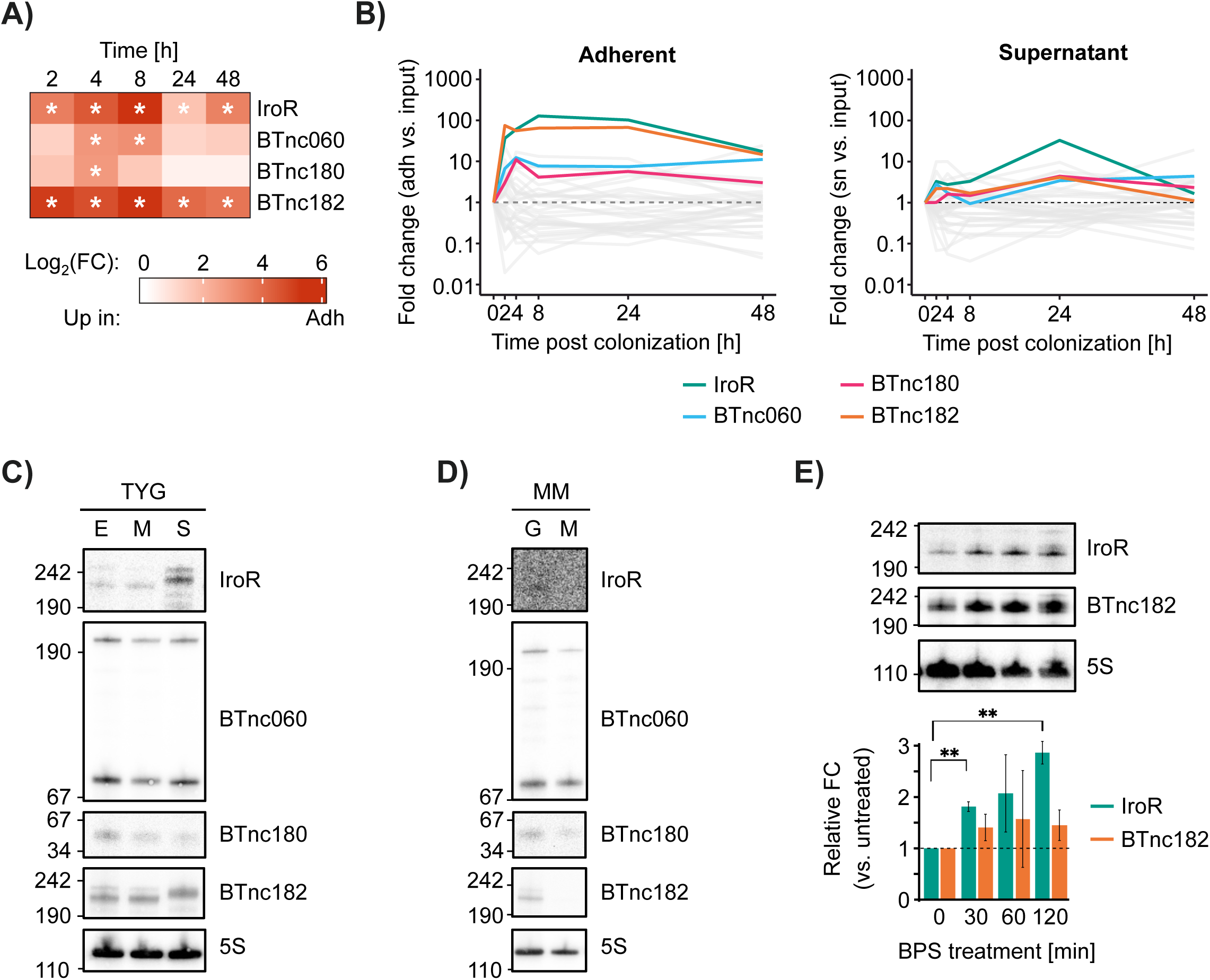
Mucosal-specific *Bacteroides* sRNA expression. A) Expression of selected sRNAs in adherent versus supernatant bacteria. An asterisk indicates a FDR < 0.05. B) Expression of sRNAs in adherent and supernatant bacteria relative to the input culture (time point 0). Only sRNAs with FDR < 0.05 in at least one time point are shown. IroR, BTnc060, BTnc180, and BTnc182 are highlighted. C) Northern blot validation of IroR, BTnc060, BTnc180 and BTnc182 cultured in rich (TYG) medium. E: early exponential phase, M: middle exponential phase, S: stationary phase. The 5S rRNA is shown as a loading control. D) Northen blot of IroR, BTnc060, BTnc180, and BTnc182 cultured in minimal media. G: glucose, M: mucin as sole carbon source, grown to middle exponential phase. The 5S rRNA is shown as a loading control. E) Northern blot of IroR and BTnc182 in *B. thetaiotaomicron* cultured in BHI media and grown to early exponential phase. Iron limitation was induced with 200 µM BPS and RNA extracted 0 (untreated), 30, 60, and 120 minutes later. The 5S rRNA is shown as a loading control and asterisk indicates significance with Students T-test.

Since these sRNAs are predicted from RNA-seq datasets (**Suppl. Fig. S4**) [28, 29], but have remained uncharacterized, we proceeded to validate their expression by northern blot, which revealed a number of isoforms for several of the sRNAs (**Fig. 4C**). In particular, the expression patterns of IroR, BTnc060, and BTnc182 were strongly growth phase-dependent in TYG medium. To further characterize their regulation, we assessed sRNA expression in minimal media containing either glucose (as in DMEM) or mucin as the sole carbon source (**Fig. 4D**). Notably, IroR and BTnc182 were detected in glucose, but not mucin, suggesting that expression of these sRNAs in the adherent population could be stimulated by factors other than the presence of mucus glycans alone. As the above data implied the mucosal niche to be depleted of iron and since sRNAs are key regulators of iron homeostasis across diverse bacteria [87], we hypothesized that IroR and BTnc182 may instead respond to low iron concentrations. To test this, we cultured *B. thetaiotaomicron* to early exponential phase in brain heart infusion (BHI) media, then induced iron-limitation with the iron chelator bathophenanthroline disulfonic acid (BPS). While expression of BTnc182 was not significantly altered during iron limitation, IroR was significantly induced 30 minutes after BPS treatment and levels were maintained 60 and 120 minutes later (**Fig. 4E**), suggesting that IroR could have a role in iron homeostasis.

### IroR is a conserved Bacteroides sRNA involved in iron homeostasis

In contrast to the intestinal lumen, the gut mucous layer is dominated by host-derived glycans as the primary carbon source for resident commensals but is markedly depleted in the essential micronutrient iron [14]. As nearly all organisms require iron as a cofactor for essential cellular processes, including energy generation and DNA replication [88], this scarcity poses a challenge for bacterial colonizers. Indeed, in murine models of gut epithelial colonization, iron acquisition dominates the transcriptional response of *E. coli* in the colonic mucus, surpassing even oxygen and carbon source availability [14]. Since the hypoxic growth conditions of our Transwell model likely recapitulate the iron-limiting environment at the mucosal epithelium, we hypothesized that this iron deprivation could be a major driver of the observed transcriptomic remodeling. We therefore performed an RNA-seq experiment on *B. thetaiotaomicron* cultured in BHI media (iron-replete) or BHI supplemented with BPS (iron-limiting). Consistent with the above data, expression of IroR was robustly upregulated when *B. thetaiotaomicron* was iron-starved (**Fig. 5A**). Sequence analysis shows that IroR is conserved amongst *Bacteroides* spp. (**Suppl. Fig. S5A**) and OrfFinder found no open reading frames within the sRNA [89], supporting its annotation as noncoding. Structural prediction using RNAfold [90] suggests that IroR would form features characteristic of regulatory sRNAs, including an extensive bulged 5′ hairpin that is followed by a stem loop and an intrinsic terminator (**Suppl. Fig. S5B**).

**Figure 5:**
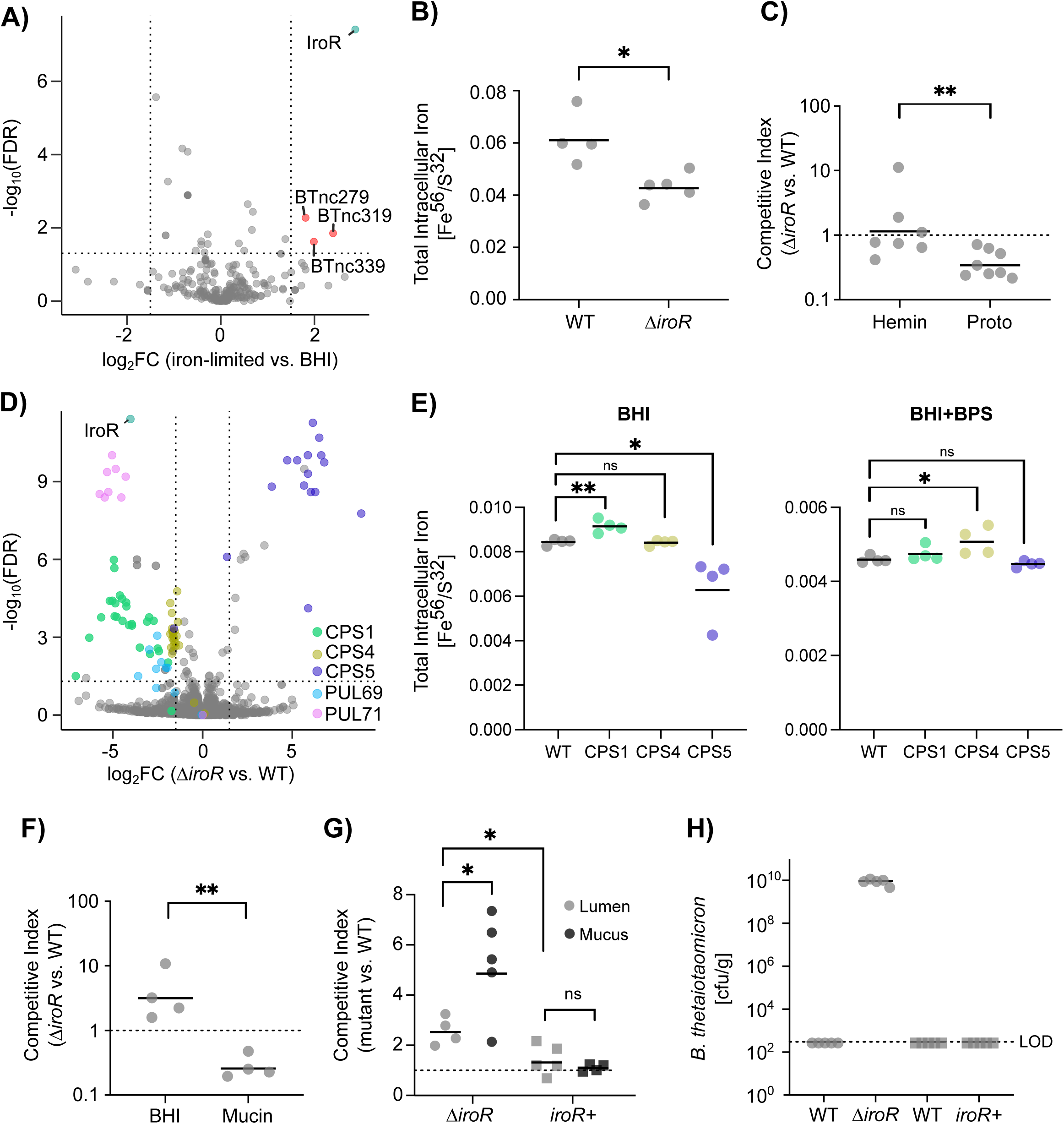
IroR contributes *B. thetaiotaomicron* iron homeostasis *in vitro* and *in vivo*. A) Volcano plot showing differential expression of *B. thetaiotaomicron* sRNAs in iron-limited (BHI + 200 µM BPS) vs. mock-treated (BHI) cultures. B) The intracellular iron content of wild-type *B. thetaiotaomicron* and Δ*iroR* in BHI media supplemented with 200 µM BPS. Quantified by ICP-MS. Asterisk indicates significance with Students T-test. C) Competitive index for wild-type *B. thetaiotaomicron* vs. Δ*iroR* grown in semi-defined media (SDM) supplemented with hemin (iron-replete) or SDM supplemented with protoporphyrin (iron-limited). D) Volcano plot showing differential gene expression of wild-type *B. thetaiotaomicron* compared to Δ*iroR,* done 15 minutes after treatment with BPS. E) The intracellular iron content of wild-type *B. thetaiotaomicron* and mutants expressing a single CPS locus in BHI media (left) or BHI media supplemented with 200 µM BPS. Quantified by ICP-MS. Asterisk indicates significance with Students T-test. F) Competitive index for wild-type *B. thetaiotaomicron* vs. the Δ*iroR* in BHIS broth or porcine mucin broth. G) Competitive index for *iroR* deletion or complementation mutants (Δ*iroR* and *iroR*+) vs. WT in the cecal contents (lumen or mucosal layer) of colonized mice six days post-inoculation. Asterisk indicates significance with Students T-test. H) Competitive colonization for *iroR* mutants vs. WT in the luminal contents of colonized mice nine days post-inoculation.

To test whether IroR plays a role in regulating iron homeostasis in *B. thetaiotaomicron*, we generated a non-polar deletion mutant strain (Δ*iroR*) and assessed its intracellular iron content using inductively coupled plasma mass spectrometry (ICP-MS). Consistent with our hypothesis, the deletion mutant exhibited a significant reduction of total intracellular iron compared to the wild-type when extracellular iron was limited (**Fig. 5B**). We next examined whether this defect in iron homeostasis impairs *B. thetaiotaomicron* fitness by competing the wild-type and Δ*iroR* in a semi-defined medium (SDM) supplemented with hemin (iron-replete) or the protoporphyrin cofactor lacking iron (iron-limited). This experimental design allowed us to modulate iron availability while maintaining tight control of other nutritional variables in the culture media. The wild-type significantly outcompeted Δ*iroR* in iron-limited SDM, but not under iron-replete conditions (**Fig. 5C**). Together, these data indicate that the sRNA IroR is induced in response to iron-starvation *in vitro* and during colonization of the mucous layer of the Transwell model, and that it contributes to *B. thetaiotaomicron* iron homeostasis and iron-dependent fitness.

### IroR modulates B. thetaiotaomicron capsule gene expression

To elucidate the mechanisms by which IroR regulates iron homeostasis in *B. thetaiotaomicron*, we compared the transcriptomes of the wild-type and Δ*iroR* following iron restriction. Cultures were grown to early exponential phase in BHI media, then treated with the iron-chelator BPS to induce IroR expression, and subjected to RNA sequencing 15 minutes post-treatment. Notably, in comparison to the treated wild-type strain, the majority of differentially expressed genes in the Δ*iroR* mutant were associated with CPS loci (**Fig. 5D**). Among the eight known CPS genomic loci in *B. thetaiotaomicron*, CPS5 transcripts were significantly increased in Δ*iroR* compared to wild-type, suggesting that IroR represses CPS5 expression under iron-limited conditions. In contrast, CPS1 and CPS4 transcripts were significantly lower compared to the wild-type.

Given this differential expression of CPS loci, we tested whether specific capsule types differ in their capacity to support iron uptake. Using ICP-MS we compared the intracellular iron content of wild-type *B. thetaiotaomicron* and mutant strains expressing only a single capsule type; CPS1, CPS4, or CPS5. In BHI alone, the CPS1-expressing strain had a significantly higher intracellular iron content than wild-type, while the iron content of the CPS5-expressing strain was significantly lower (**Fig. 5E**). In contrast, when iron-limitation was induced with BPS, the CPS4-expressing strain exhibited a significantly higher iron content compared to the wild-type. These results support the hypothesis that IroR promotes iron acquisition by modulating the expression of capsule types, increasing CPS1 and CPS4 which are more permissive to iron capture and decreasing CPS5.

In addition to capsule regulation, RNA-seq during iron-limitation suggests IroR upregulates PULs 69 and 71 (**Fig. 5D**), whose putative substrates are alpha-mannan and mucin *O*-linked host glycans, respectively. Therefore, IroR could be facilitating a shift in glycan preference toward host-derived polysaccharides when iron is limited, in line with our hypothesis that iron availability is a major driver of *B. thetaiotaomicron* adaptation to the mucous layer niche. Furthermore, wild-type *B. thetaiotaomicron* significantly outcompeted Δ*iroR in vitro* when mucin was the primary carbon source, but crucially not in BHI, in which peptides are the main carbon source (**Fig. 5F**).

### IroR influences B. thetaiotaomicron fitness in the murine gut

Having established a role for IroR in regulating adaptation to mucin utilization and iron limitation *in vitro*, we sought to assess its contribution to colonization of the intestinal mucous layer *in vivo*. Groups of C57BL/6 mice were treated with an antibiotic cocktail to permit engraftment, as conventionally raised mice are resistant to *B. thetaiotaomicron* colonization [91]. The mice were then inoculated with an equal ratio of *B. thetaiotaomicron* wild-type and the Δ*iroR* mutant. Upon inoculation, animals were switched to a low-fiber diet to reduce the availability of microbiota-accessible dietary glycans and promote foraging of host-derived mucosal glycans [10, 92, 93], thereby allowing us to model the mucous layer niche. The abundance of each strain in the luminal and cecal mucosal contents was determined six days post-inoculation by plating on selective media. Notably, the wild-type was outcompeted by Δ*iroR* in the cecal lumen, and this competitive advantage was even greater in the mucous layer (**Fig. 5G**). *Trans*-complementation of IroR (strain *iroR*+) ablated the competitive advantage, demonstrating the fitness benefit is specific to IroR (**Fig. 5G**). Moreover, the fitness benefit of Δ*iroR* was even more pronounced upon longer colonization. After nine days of co-colonization, the wild-type could not be recovered in the cecal lumen above the limit of detection, whereas Δ*iroR* bacteria were present in high numbers (∼10^10^ CFUs/g feces) (**Fig. 5H**). In mice co-colonized with the wild-type and the *iroR*+ complementation, both strains were below the limit of detection at this time point (**Fig. 5H**). Taken together, our findings suggest the fitness contribution of IroR is context-dependent, differentially influencing *B. thetaiotaomicron* colonization dynamics across gut microenvironments.

## DISCUSSION

In this study, we established a novel, tractable model to study host-bacterium interactions under hypoxic conditions and used dual RNA-seq to investigate transcriptomic changes during host colonization by an anaerobic commensal bacterium. *B. thetaiotaomicron* successfully colonized the HT29 Transwell model, forming two distinct subpopulations (**Fig. 1C**), including an adherent population residing within the mucous layer. Despite this close interaction with bacteria, the integrity of the epithelial barrier remained intact for up to 48 hours (**Fig. 1D**), with no notable increase in cytotoxicity compared to mock colonization (**Fig. 1E**). Accordingly, the host mounted a delayed and muted transcriptional response to *B. thetaiotaomicron* compared to pathogenic species such as *S*. Typhimurium [61]. It was only at 24 and 48 h p.c. that we could detect transcriptomic changes, as host cells upregulated pathways involved in ER stress (**Fig. 2B**), cytoskeletal rearrangement, cell adhesion, and cell-cell junctions (**Fig. 2C**). We hypothesize that prolonged exposure to *B. thetaiotaomicron* prompts epithelial barrier tightening, thereby preserving barrier integrity and, *in vivo*, preventing bacterial translocation into deeper tissues. This observation aligns with previous findings showing that colonization by various probiotic species strengthens the gut epithelial barrier [94–96]. Interestingly, epithelial cells respond to colonization by both *B. thetaiotaomicron* (this study) and *S*. Typhimurium [61] by repressing the let-7 miRNA family. Let-7 repression at late stages of *B. thetaiotaomicron* colonization may contribute to the host tolerance of beneficial commensal bacteria, while being primed to mount an immune response in case of pathogenic stimuli.

This dual RNA-seq approach also allowed us to distinguish transcriptional heterogeneity between the two spatial bacterial subpopulations within the Transwell model. Adherent bacteria residing within the mucous layer upregulated pathways involved in carbon, energy, and amino acid metabolism throughout the time-course, indicating a metabolically active state (**Fig. 3B-i**). We noted that several of these processes are iron dependent, and since the mucous layer is known to be an iron-limiting environment, we quantified the expression of genes involved in iron acquisition and utilization. For example, up-regulation of *xusB*, required for xenosiderophore capture [97], in the adherent subpopulation suggests there is an iron-sparing response (**Fig. 3C-i**). Additionally, several PULs, mostly associated with the consumption of host-derived glycans, were upregulated within the mucosal niche (**Fig. 3B-ii**). This is consistent with previous transcriptomic studies in mono-colonized mice, which showed that mucus-associated *B. thetaiotaomicron* differentially expressed PULs targeting host-derived polysaccharides compared to the luminal populations [21, 22]. The Transwell model, therefore, effectively recapitulates this distinct mucus-associated niche, characterized by the elevated expression of PULs associated with host-derived polysaccharides.

The expression of microbial surface structures such as pili, adhesins, and capsular polysaccharides constitutes vital components for the interactions between bacteria and their hosts [83][98]. In particular, CPS8 has been shown to be more highly expressed in luminal *B. thetaiotaomicron*, which was linked to biofilm formation on food particles [14]. Although not reaching statistical significance, CPS8 tended to be more highly expressed in the supernatant population of the Transwell model (**Fig. 3B-iii**). Another capsule type, CPS1, is strongly repressed in *B. thetaiotaomicron* biofilms [99]. Accordingly, we observed significantly higher expression of CPS1 in the supernatant subpopulation (**Fig. 3B-iii**), likely due to the lack of suitable substrates for biofilm formation in this environment. Instead, repression of this capsular locus could promote attachment to the host mucus. Indeed, a strain that constitutively expresses CPS1 (CPS1 lock; [81]) was significantly outcompeted by its wild-type counterpart in the Transwell model (**Fig. 3D**), demonstrating the importance of bacterial cell surface modulation for host-colonization. In this context, it will be interesting to test whether strains lacking selected CPS loci adhere to the host mucus more tightly or, whether strains restricted to a single CPS type [81] exhibit reduced attachment efficiency compared to wild-type. Given the well-documented challenges in correlating adhesion propensity observed *in vitro* with *in vivo* colonization capacity [83, 100, 101], our Transwell model provides a physiologically relevant platform to test *Bacteroides* mucus adherence in a simplified context free of confounding factors like host immunity and peristaltic flow.

Well-studied model bacteria, e.g. species belonging to the *Pseudomonadota* (formerly *Proteobacteria*), employ regulatory RNAs to shape their interaction with host cells and tissues [27]; however, whether this post-transcriptional gene expression control extends to *Bacteroides* spp. has not previously been explored [86]. We therefore directed our attention to sRNAs that were upregulated in adherent bacteria as these could potentially be important factors driving the adaptation of *B. thetaiotaomicron* to this gastrointestinal niche. Among these, IroR was consistently expressed at high levels throughout colonization of the Transwell model (**Fig. 4AB**). Interestingly, we were unable to detect this sRNA during *in vitro* growth where mucin served as the sole carbon source (**Fig. 4D**). Since the adherent population may experience iron-limitation (**Fig. 3C-i**), we hypothesized that iron could be a potential cue driving IroR expression. We compared the transcriptome from *B. thetaiotaomicron* grown in iron-limited media and found that IroR was highly expressed, supporting our hypothesis that its induction in the adherent subpopulation is triggered by iron limitation, rather than the presence of mucins (**Fig. 5A**).

Given the reliance of numerous essential enzymes on iron as an essential cofactor, iron homeostasis is tightly regulated in bacterial cells. In *Enterobacteriaceae*, the conserved transcriptional repressor Fur complexes with ferrous iron and binds to its target promoters, suppressing genes that encode iron uptake proteins, iron-independent proteins, and the sRNA RyhB [102]. Upon iron starvation, apo-Fur disassociates from its target promoters, licensing the transcription of its regulatory targets, including RyhB [103]. Via imperfect complementary base-pairing and stabilization by the RNA chaperone Hfq, RyhB binds to the mRNA transcripts of iron-rich proteins, blocking their ribosome-binding site or targeting them for degradation [35, 104, 105]. Since the pioneering studies on *E. coli* RyhB [35, 106–108], several sRNAs in diverse species have been identified as post-transcriptional regulators that facilitate adaptation to iron limitation [109–120]. However, in *Bacteroidota* neither a Fur homolog, nor a regulatory RNA component of iron homeostasis control has been previously reported.

Further investigation into the functional role of IroR suggested it regulates the expression of CPS loci (**Fig. 5D**), whose expression was differentially regulated between the adherent and supernatant subpopulations. Mechanistically, it appears unlikely that a CPS-derived transcript is a direct target of IroR, as *in silico* analyses did not reveal regions of extended sequence complementarity with this sRNA. Notably, CPS1 was upregulated by IroR under iron-limited conditions, contrasting with our findings from the dual RNA-seq experiment. This discrepancy suggests that additional negative regulators may override IroR-dependent upregulation of CPS1 repression in mucus-adherent bacteria in the presence of host mucus. Further investigation is needed to elucidate the complexities of this regulatory network.

Our *in vivo* observation that Δ*iroR* outcompetes the wild-type in the mucous layer (**Fig. 5G**) raises the question as to why *B. thetaiotaomicron* would maintain a factor that is disadvantageous to its niche colonization and fitness. In the Transwell model, *B. thetaiotaomicron* repressed expression of CPS1 and CPS4 in the adherent fraction, suggesting these capsule types may reduce fitness in the mucous layer niche. This is supported by the significant colonization fitness defect of the CPS1-locked mutant (**Fig. 3F**). However, despite their apparent fitness cost, CPS1 and CPS4 were up-regulated by IroR during iron limitation *in vitro*. As mutants expressing only CPS1 or CPS4 capsule types had higher intracellular iron levels than the wild-type strain (**Fig. 5E**), IroR may mediate a tradeoff between capsule-mediated iron acquisition and mucus colonization fitness. Thus, we suggest IroR functions to fine-tune capsule expression in response to environmental cues, balancing iron acquisition with niche-specific fitness demands in a manner that supports adaptation across diverse gut conditions.

## SUPPLEMENTARY FILES

**Figure S1:**
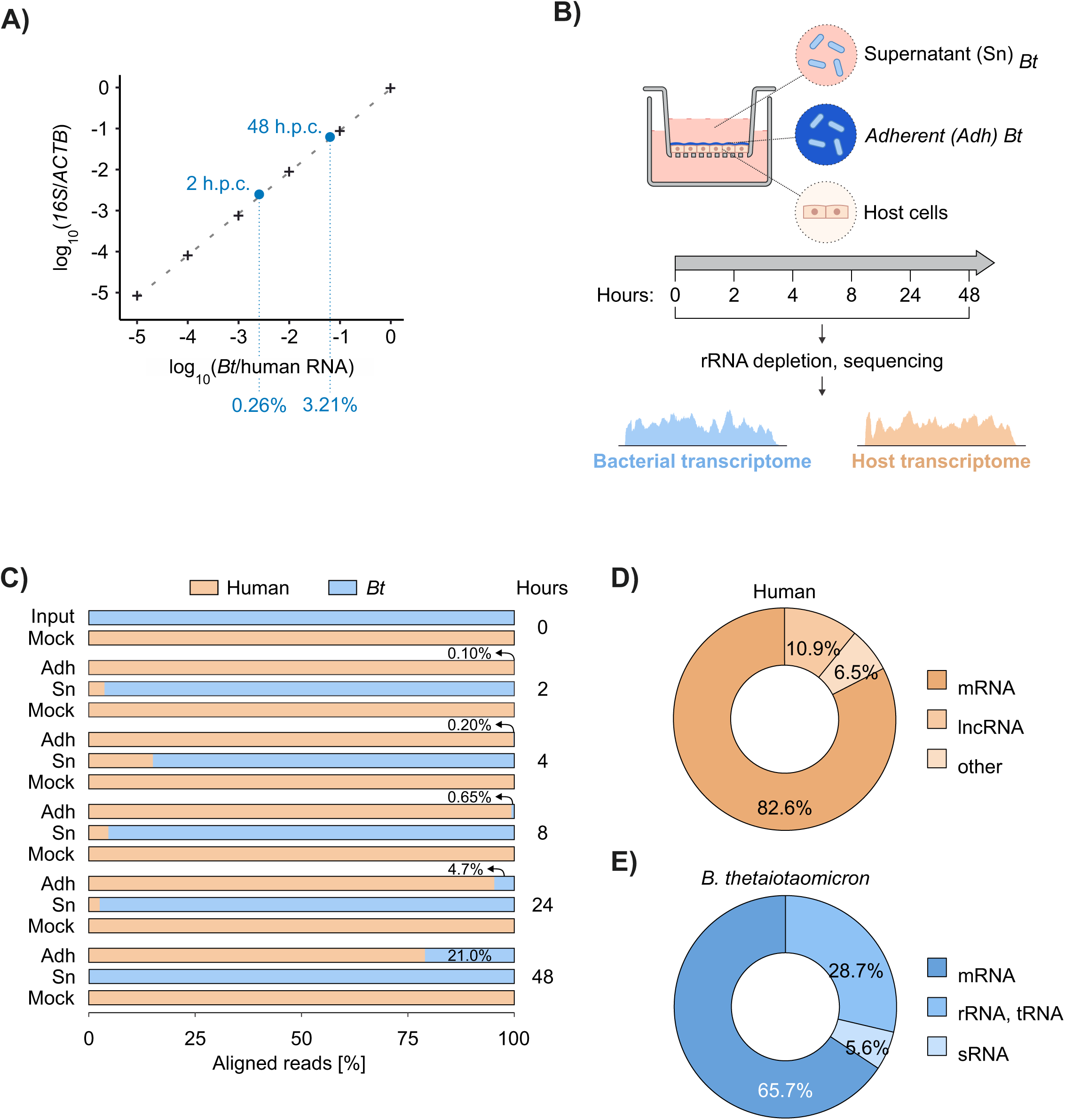
A) qRT-PCR-based determination of bacterial/host RNA ratio. The gray dashed line represents the standard curve fitted to qPCR values (black crosses) of mixtures of a constant amount of human RNA and progressive dilutions of *B. thetaiotaomicron* total RNA. The x-axis represents the mix ratio of bacterial to human RNA and the y-axis is the ratio between the resulting qPCR Ct values of the *Bacteroides* 16S rRNA and human *ACTB* genes. Data points for RNA extracted from colonized Transwells are plotted in blue and their interpolated value is shown below. B) Experimental workflow. At the specified timepoints, host cells, supernatant and adherent *B. thetaiotaomicron* are collected, RNA extracted and sequenced. This provides a snapshot of the host and bacterial transcriptomes over the course of colonization. C) Mapping statistics of representative samples, indicating the fraction of reads mapping to the human or *B. thetaiotaomicron* transcriptomes. D) RNA class distribution in a representative human sample. E) RNA class distribution in a representative bacterial sample.

**Figure S2:**
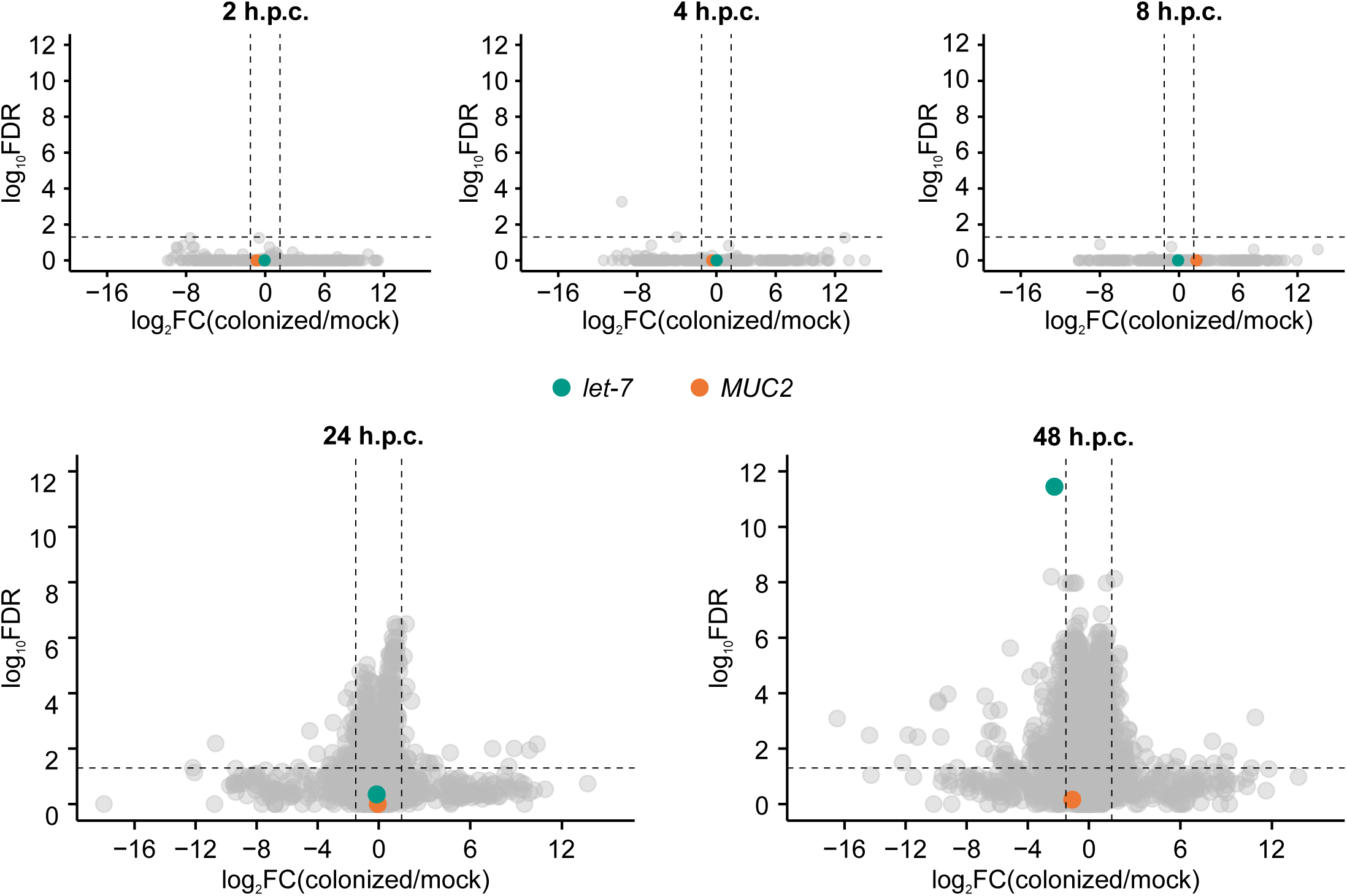
Volcano plots of fold change of colonized versus mock host cells and false discovery rate at all time points. Vertical dashed lines mark -1.5 and 1.5 log_2_FC, while the horizontal line indicates the 0.05 significance threshold. The *MUC2* and *MIRLET7A1HG* genes are highlighted in colors.

**Figure S3:**
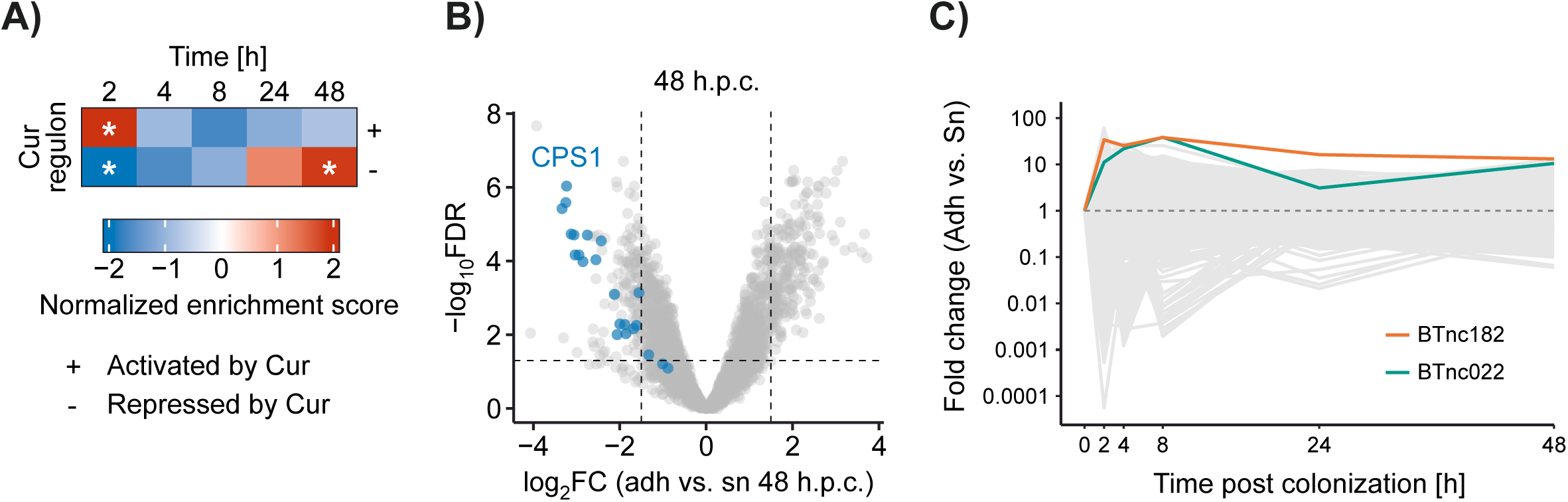
A) Enrichment of the Cur regulon between adherent and supernatant bacteria. NES: normalized enrichment score. An asterisk indicates FDR < 0.05. B) Volcano plot showing RNA-seq data from adherent versus supernatant bacteria at 48 h p.c. Genes belonging to CPS1 are colored in blue. C) Plot of all bacterial genes over the time course of colonization. Expression of sRNAs in adherent versus supernatant bacteria. Only genes with FDR < 0.05 in at least one time point are shown.

**Figure S4:**
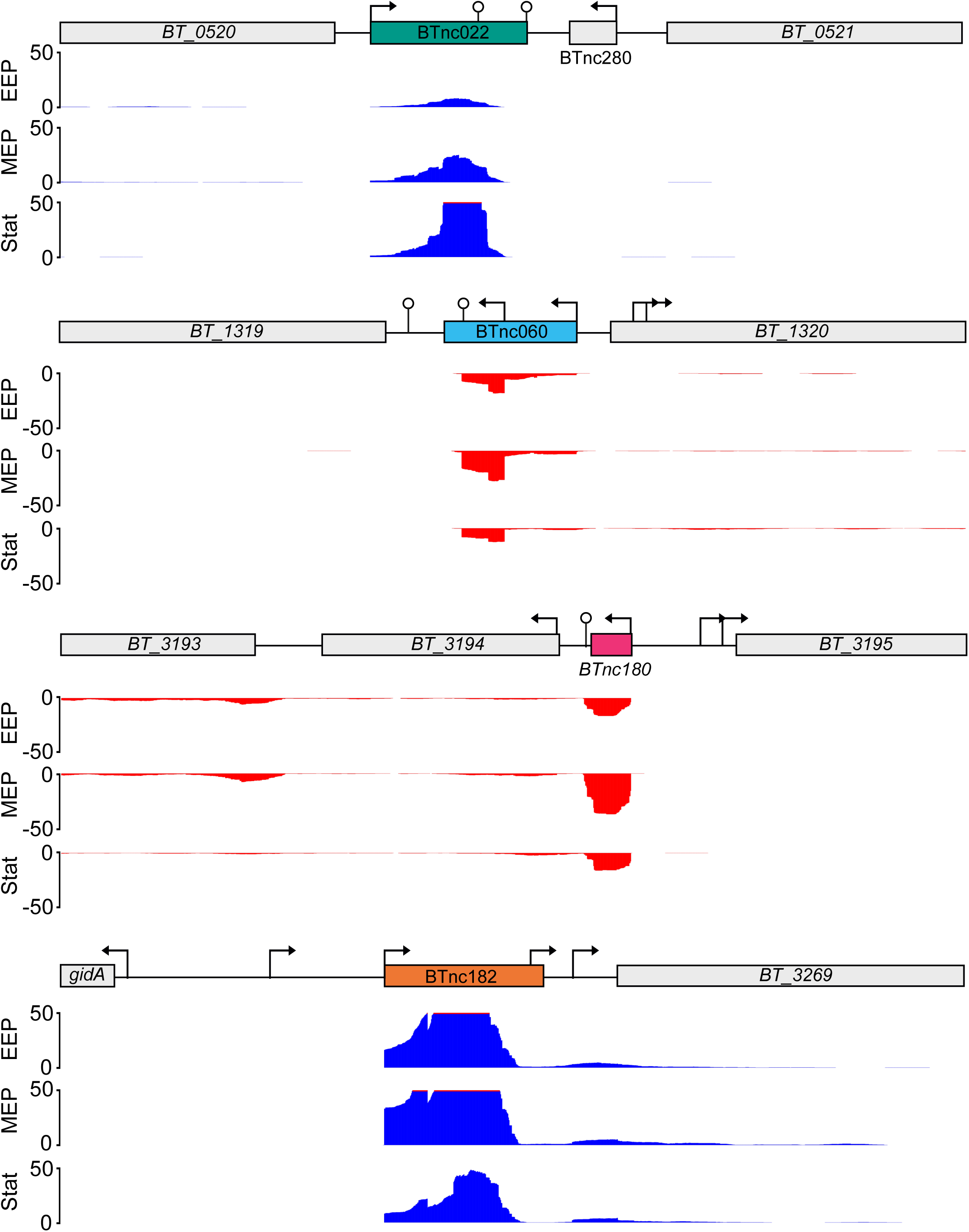
RNA sequencing read coverage for IroR, BTnc060, BTnc180 and BTnc182. A 1.5 kb window is shown with adjacent genes, start site and terminator annotations from *B. thetatiotaomicon* cultures grown in TYG [29].

**Figure S5:**
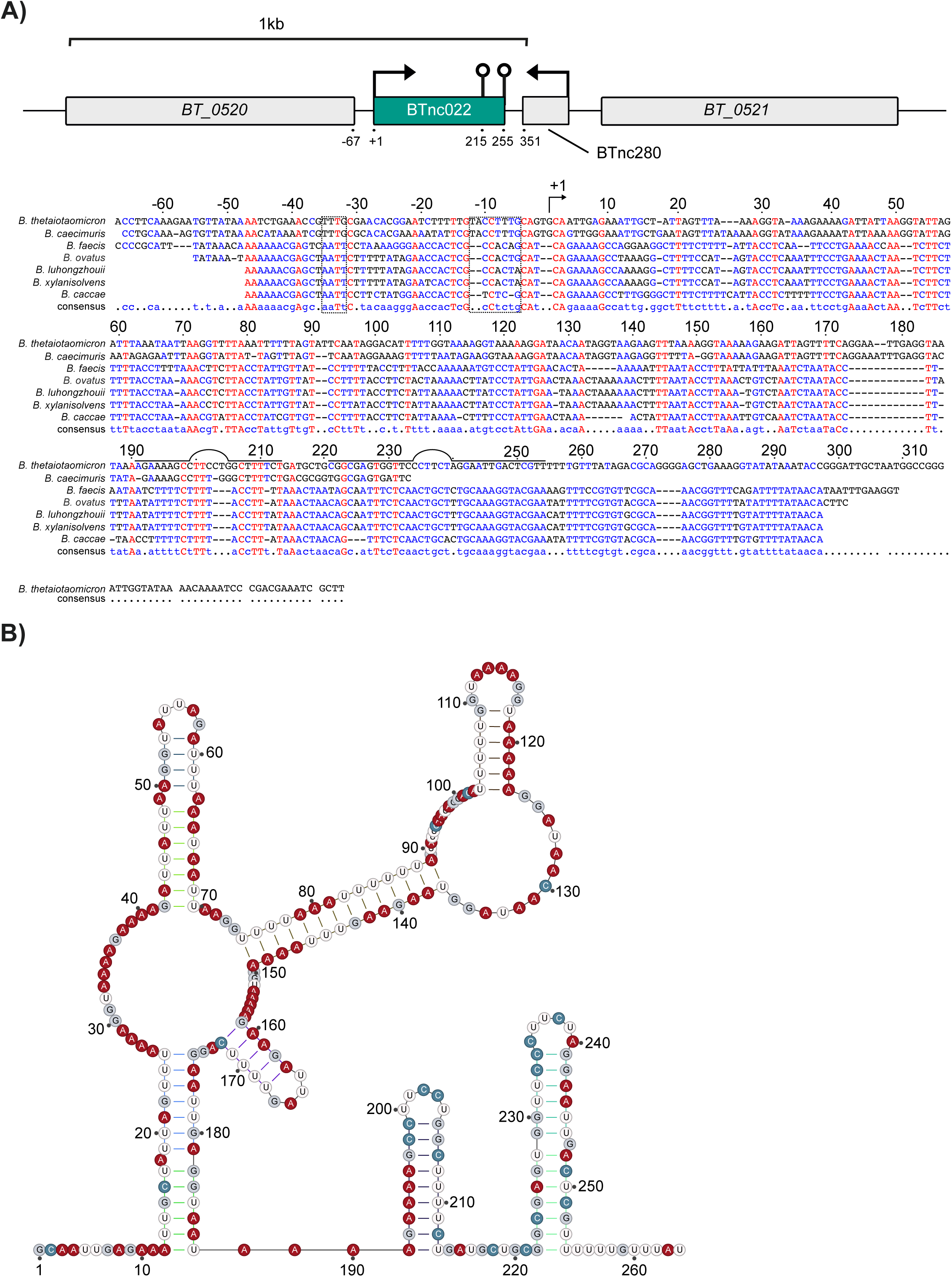
A) Genomic context and conservation of IroR amongst *Bacteroidetes* at the nucleotide level. Red and blue colors indicate conserved and less-conserved ribonucleobases, respectively. Numbers denote position relative to the 5′ end of IroR. Predicted -10 and -35 promoter elements are boxed. B) Predicted structure of IroR.

## MATERIALS AND METHODS

### Bacterial strains and genetics

A list of strains, oligonucleotides, and media used in this study can be found in Table 1. Overnight cultures (∼16 h) of *Bacteroides thetaiotaomicron* strain VPI-5482 were cultured in TYG or BHI medium at 37°C in an anaerobic chamber (Coy Laboratory Products) using an anaerobic gas mix of 85% N2, 10% CO2, and 5% H2. To construct deletion mutants the pExchange-tdk or pSIE1 plasmid systems were used as previously described [122, 123]. In brief, flanking regions of the gene of interest were cloned into either suicide plasmid, conjugated into *B. thetaiotaomicron* using *E. coli* S17 λ-pir as the donor strain and selected using the appropriate antibiotics. Following counter selection, colonies were confirmed by PCR and sanger sequencing. For complementation of IroR *in-trans* using the pNBU2 plasmid [124], IroR was amplified and expressed from its native promoter.

### Human cell line and culture conditions

The HT29-MTX-E12 cell line was a gift of Prof. Cynthia Sharma. Cells were routinely cultured in 5% CO_2_ and atmospheric O_2_ in DMEM medium supplemented with 10% fetal bovine serum, 1% (v/v) L-glutamine, 1% (v/v) non-essential amino acids and 1% (v/v) sodium pyruvate. Cultures were split at ∼80% confluence and maintained for up to 20 passages.

Gut epithelial models were prepared as follows. 12-well Transwell inserts (Transwell, Corning) were seeded with 4x10^5^ cells in 0.5 mL medium added to the apical side and incubated 20 min at 37°C, after which 1.5 mL medium were added to the basolateral compartment and the plate was returned to the incubator. After 2 days, media in both compartments was replaced with the same volumes and the plate was moved in the incubator on an orbital shaker set at constant 65 rpm speed. Media was replaced every 2-3 days, until complete model maturation at 21-28 days post-seeding.

### Histology and immunofluorescence staining of Transwell models

After removal of basolateral and apical medium, the models were washed once with PBS in both compartments. Cells were then fixed in Methacarn solution (60% methanol, 30% chloroform, 10% glacial acetic acid) overnight at 4 °C. After this, the fixed cell layer was incubated twice in 100% methanol, followed by twice in 100% ethanol, for 10 minutes each. The Transwell membrane with adherent, fixed cells was then carefully excised with a scalpel, wrapped in methanol-soaked embedding paper and inserted in an embedding cassette. These were then placed in an embedding machine (Thermo Fisher, A81000002), on which an automated protocol was run, composed of 2x incubation in 100% xylene for 15 minutes, followed by 2x in paraffin for 60. Next, cassettes were incubated at 60 °C for 20 minutes to remove excess paraffin. Samples were then extracted from each cassette, cut into half and blocked in stainless steel casting mold with paraffin. Blocks were then cut into 5 μm slices and dried at 37 °C overnight. Dried slices were then incubated at 60 °C for one hour to melt the paraffin and incubated in the following solutions for deparaffinization and rehydration: 2x 100% xylene for 10 minutes, dipping 3x in 96 % ethanol and 3x in a second, fresh 96 % ethanol aliquot, followed by 3x dipping in 70% ethanol, 3x dipping in 50% ethanol and finally swirling in deionized water until clear. Hematoxylin-eosin staining was then performed by incubating samples as follows: 6 minutes in hematoxylin (Morphisto, 10231) rinsing in deionized water until clear, dipping in HCl and water, 5 minutes in tap water, 6 minutes in eosin (Morphisto, 10177) and finally in deionized water until clearing. Staining of mucus was then done by incubation for 3 minutes in 3% acetic acid, 15 minutes in 1% alcian blue (Morphisto, 10126), rinsing in deionized water until clear, 5 minutes in Nuclear Fast Red stain (Morphisto, 10264) and finally in deionized water until clear. Samples were then dehydrated by dipping twice in 70% ethanol, incubation for 2 minutes in 96% ethanol, 5 minutes in isopropanol, 5 minutes in fresh isopropanol, 5 minutes in xylene and finally 5 minutes in fresh xylene. Stained slices were then mounted on SuperFrost®Plus glass slides (Langenbrinck, 03-0063) with Entellan (Merck, 1079600500) medium. Imaging was done on an inverse fluorescence microscope (Keyence, BZ-9000).

For immunofluorescence analyses, fixed membranes were permeabilized in 0.2% Triton X-100 in PBS and washed three times with PBS-0.5% Tween (PBS-T). In order to minimize non-specific antibody binding, the sample underwent a 30-min incubation at room temperature with a 5% donkey serum solution (Abcam, ab7475) in PBS-T. Following this, primary antibodies against ZO-1 or MUC-2, diluted in PBS-T, were applied to the sample and left to incubate overnight at 4°C. Following 2x PBS-T washes the next day, secondary antibodies diluted in PBS-T were added on the membrane to incubate for 2 h at room temperature. Following a single wash in PBS-T, the samples were subjected to a 20-min incubation at room temperature with Phalloidin and/or DAPI, both diluted in PBS-T. Two subsequent PBS-T washes were performed before the samples were embedded in Fluoromount G (Invitrogen, 00-4958-02). Imaging was performed with a confocal microscope (Stellaris, Leica). The images were processed using Fiji (version 2.14.0).

### Colonization of Transwell models with B. thetaiotaomicron

On day -1, the plate containing the Transwell models was removed from the shaker and put in the incubator under static conditions. On the morning of the day of colonization, media of the Transwell models was replaced with fresh one, by adding 1.5 mL to the basolateral compartment and 400 μL to the apical one. The plate was then returned to the incubator. Next, 1 mL of an overnight *B. thetaiotaomicron* culture was pelleted, washed with 1 mL sterile PBS and centrifuged again. Cells were then resuspended in 5 mL supplemented DMEM and the OD concentration was determined. This cell suspension is the input culture used for colonization. Transwell models were removed from the incubator and 4x10^6^ bacterial cells (MOC 10) resuspended in 100 μL supplemented DMEM were added to the apical compartment. After centrifugation at 250g for 10 minutes, plates were moved to a hypoxic incubator (1% pO_2_, 5% CO_2_, 37 °C) and kept undisturbed for up to 48 hours.

### CFU, permeability, and cytotoxicity assays

For CFU assays, 50 μL of the supernatant medium was collected, then the remaining DMEM was removed and the epithelial layer was washed once with PBS. Host cells, mucus and adherent bacteria were at this point collected in PBS by scraping with a pipette tip and moved to a 2 mL tube. Samples were homogenized by repeated pipetting. Supernatant and adherent samples were then serially diluted in PBS and plated on BHIS agar plates in 3x10 μL spots. Plates were incubated anaerobically at 37°C for two days, after which spots with 20 to 200 colonies were enumerated.

Permeability (tightness) of the host cell layer was measured as follows. FITC-dextran (Sigma, G2838) was dissolved in supplemented DMEM at a concentration of 0.25 mg/mL and added to the apical compartment. The model was then returned in the incubator for 30 min, after which the fluorescent emission of 3x100 μL samples of each basolateral compartment was measured on a Tecan microplate reader. Values were normalized to the intensity of the original FITC-dextran DMEM suspension. A value below 1% was considered indication of sufficient epithelial barrier tightness.

For host cytotoxicity measurements, the activity of cytosolic lactate dehydrogenase (LDH) in the supernatant medium was measured with the CytoTox 96® Non-Radioactive Cytotoxicity Assay kit (Promega, G1780) according to the manufacturer’s protocol. Briefly, 50 μL of the supernatant medium were pipetted in a well of a 96-well plate and mixed with an equal volume of reconstituted Substrate Mix. Samples were then incubated at room temperature for 30 min in darkness, after which 50 μL of Stop solution were added to each well. Light absorption at 490 nm was then measured on a Tecan Microplate Reader. Data were normalized to those of time point 0 models (set to 1).

### Determination of the bacterial-to-host RNA ratio

RNA was extracted from a confluent HT29-MTX-E12 flask using Trizol reagent (Invitrogen #15596026) according to the manufacturer’s protocol and an early exponential phase (OD 0.5) *B. thetaiotaomicron* culture with the hot phenol method as described previously [125]. Contaminating DNA was removed by digestion of 40 μg of RNA with 5 U of DNase I (Thermo Fisher Scientific #EN0521), followed by purification with a phenol-chloroform extraction followed by ethanol-NaOAc precipitation.

Estimation of total RNA ratios was done as previously described [48, 125]. Briefly, serial 1:10 dilutions of *B. thetaiotaomicron* total RNA were mixed with a constant quantity (0.5 ng) of human RNA as indicated and the amount of each quantifiead through one-step RT-qPCR of the human *ACTB* and bacterial 16S rRNA genes. Fold changes relative to the 1:1 mixture were then calculated and a linear regression line was fit to the resulting data points. The respective value of colonization samples was then interpolated into the line to obtain the corresponding bacterial to host RNA ratio.

### Dual RNA-seq: RNA extraction, library preparation, and sequencing

Dual RNA-seq samples were collected in biological triplicates. At the moment of sample harvesting, 400 µL of supernatant medium were slowly collected from the Transwell models and moved to individual 2 mL tubes. The remaining medium was then aspirated and the apical portion of the Transwell washed once with 500 µL PBS. Next, 500 µL Trizol reagent was added to the apical compartment and set aside for at least 5 minutes. In the meantime, 1.2 mL Trizol LS reagent (Invitrogen #10296028) was mixed into the supernatant sample. After incubation of the Transwells with Trizol, the cell material was resuspended and detached with the help of a pipette tip with care not to pierce the Transwell membrane and transferred to a 1.5 mL tube. At this point, chloroform was added to the supernatant and adherent tubes at a volume of 100 and 320 µL, respectively. After mixing, RNA was extracted according to the Trizol protocol and precipitated with one volume of isopropanol and 0.75 µL glycoblue (Invitrogen #AM9516). cDNA libraries were generated at the Core Unit SysMed (University of Würzburg, Germany) using the Illumina Stranded Total RNA Prep Ligation with Ribo-Zero Plus protocol according to manufacturer’s recommendation with the following modifications: for total RNA samples with 1-40 ng RNA input 15-17 PCR cycles, for samples with 100 ng total RNA input 14 PCR cycles. After quality control, libraries were sequenced on an Illumina NextSeq 500 platform (1x75 bp single-end run).

### Read processing and mapping; differential gene expression and gene set enrichment analysis

Reads were processed, aligned and quantified with the dualrnaseq nf-core pipeline (https://nf-co.re/dualrnaseq/) with options *--run_bbduk --single_end -- run_salmon_selective_alignment --libtype A*. The human genome assembly used was GRCh38.p13.

Differential gene expression analysis was done with *edgeR* (3.32.1) separately for human and bacterial reads. For the host, lowly expressed genes were filtered out with the *filterByExpr* function and the TMM method of normalization was used. For *B. thetaiotaomicron*, we kept genes with cpm >1 in >25 samples and libraries were full-quantile normalized with *EDASeq* [126]. The *withinLaneNormalization* function of the same package was used to correct for observed gene length-dependent biases observed in bacterial read counts. The *RUVs* method (k=3 for human and k=1 for bacterial libraries) was employed for batch effect correction [127].

*B. thetaiotaomicron* gene set enrichment analysis was performed on GO Terms and KEGG Pathways (both retrieved on Nov 13^th^ 2023) with the *fgsea* R package over all gene sets with ≥5 and ≤150 genes, except for PULs, which were retained independently of their gene number. Genes were ranked based on the -log_10_(PValue)*sign(fold change) metric. For human samples, we retrieved GO Terms and KEGG Pathways on Oct 21^st^ 2021 and used only gene sets with a gene number between 25 and 500.

Enrichment maps were generated on Cytoscape with the *EnrichmentMap* plugin[63] with the following cutoffs: 0.001 FDR and 0.4 combined Jaccard and Overlap coefficient (24 h p.c.), 0.00001 FDR and 0.4 combined Jaccard and Overlap coefficient. Clusters were defined with *AutoAnnotate* [121] with MCL clustering and default parameters and then manually renamed.

### Northern blot analysis

Northern blotting was performed as previously described [29]. Briefly, 5 μg or 10 µg of total RNA (extracted as described under “Determination of the bacterial/host RNA ratio“) was separated on a denaturing 6% polyacrylamide 7 M urea gel and blotted onto Hybond XL membranes (Amersham). Membranes were probed with ^32^P-labeled gene-specific oligonucleotides (Table S1).

### Iron-limitation RNA-seq

To identify iron-response sRNAs, 8 colonies of wild-type *B. thetaiotaomicron* were cultured anaerobically in 5 ml of brain-heart-infusion supplemented (BHIS) media (0.8% brain heart infusion from solid, 0.5% peptic digest of animal tissue, 1.6% pancreatic digest of casein, 0.5% sodium chloride, 0.2% glucose, 0.25% disodium hydrogen phosphate, 0.005% hemin, 0.0001% /vitamin K, pH 7.4) for 24 hours, and subcultured (1:50) in 5 ml of BHIS for an additional 5 hours, until cultures reached early exponential phase. Half of the cultures were then treated with 200 µM bathophenanthroline disulfonate (BPS) to induce iron-limitation and cultures were incubated for an additional 2 hours. Cell pellets were collected in RNAprotect (Qiagen), and RNA was extracted using TRI reagent (Molecular Research Center, Cincinnati). DNA contamination was removed using the Turbo DNA-free Kit (Ambion, USA) per the manufacturer’s recommendations. Libraries were prepared using the RNAtag-seq method [128] and sequenced on an Illumina Novaseq-SP platform (100 bp paired-end run).

### Inductively coupled plasma mass spectrometry (ICP-MS)

For measurement of intracellular iron concentration, 5 colonies of the indicated strains were cultured anaerobically in 5 ml of BHIS media for 24 hours, then subcultured (1:50) in 5 mL of BHIS for an additional 5 hours until cultures reached early exponential phase. Cultures were then treated with 200 µM bathophenanthroline disulfonate (BPS) and cultured for an additional 2 hours. Cells were washed three times with 1 mM EDTA (pH = 8.0) to remove extracellular iron before acid digestion. After collection, the samples were digested using freshly prepared 50% nitric acid (Thermo Fisher Scientific, USA). Incubation in nitric acid was allowed to proceed for two days to ensure complete digestion of the cells and dissolution of the metals to be analyzed. The samples were then diluted to 3% nitric acid. The solution was then centrifuged and filtered if needed to remove any particulates. The supernatant was analyzed for iron by inductively coupled plasma mass spectrometry (ICP-MS) using Agilent 7700x instrument (Agilent Technologies).

### In vitro competition assays in semi-defined media

For the *in vitro* competition assays, wild-type *B. thetaiotaomicron* and Δ*iroR* (marked with erythromycin resistance via integrative plasmid pNBU2-bla-ermG) strains were inoculated at (5x10^4^ CFU/mL) into a semi-defined media (1.5 g/L KH_2_PO_4_, 0.5 g/L NH_4_SO_4_, 0.9 g/L NaCl, 150 mg/L L-methionine, 5 mg/L vitamin B_12_, 20 mg/L MgCl_2_ x 6 H_2_O, 10 mg/L CaCl_2_ x 2 H_2_O, 1 mg/L MnCl_2_ x 4 H_2_O, 1 mg/L CoCl_2_ x 6 H_2_O, 1 g/L L-cysteine, 5 g/L glucose, 1 g/L tryptone, 0.2% NaHCO_3_) supplemented with 5 mg/L protoporphyrin IX or hemin. After 16 hours of competition, the cultures were selectively plated on BHI plates, treated with or without 50 µg/mL erythromycin, and incubated at 37°C for 48 h to count CFUs.

### B. thetaiotaomicron wild-type vs. ΔiroR RNA-seq

To compare the transcriptomes of the wild-type *B. thetaiotaomicron* and Δ*iroR* strains, 4 colonies of each strain were cultured anaerobically in 5 ml BHIS media for 24 hours, and subcultured (1:50) in 5 ml of BHIS for an additional 5 hours, until cultures reached early exponential phase. The cultures were then treated with 200 µM bathophenanthroline disulfonate (BPS) to induce iron-limitation and cultures were incubated for 15 min. RNA was extracted and processed for differential expression analysis as described above.

### Mouse experiments

Mouse experiments were conducted in accordance with the policies of the Institutional Animal Care and Use Committee at Vanderbilt University Medical Center (protocol # M2300019-00). C57BL/6 J wild-type (cat# 000664), were obtained from Jackson Laboratory. Mice were housed in sterile cages under specific pathogen-free conditions on a 12 h light cycle, with *ad libitum* access to food and sterile water at Vanderbilt University Medical Center. Seven to nine-week-old female or male mice were randomly assigned into treatment groups before the experiment. Antibiotic cocktails (ampicillin [Sigma-Aldrich], metronidazole [Sigma-Aldrich], vancomycin [Chem Impex International], and neomycin [Sigma-Aldrich]; 5 mg of each per mouse) were administered by oral gavage daily for 5 days. After antibiotic treatment, mice were inoculated with an equal mixture of 0.5 × 10^9^ CFU of the *B. thetaiotaomicron* wild-type strain and 0.5 × 10^9^ CFU of the respective *iroR* mutant (Δ*iroR*, *iroR*+) for 6 days. On the day of *B. thetaiotaomicron* inoculation, mice were switched to a low fiber diet (TD.86489) until the end of the experiment. After euthanasia, luminal and mucosal cecal content was collected in sterile PBS, and the abundance of *B. thetaiotaomicron* strains was quantified by plating serial-diluted intestinal contents on selective agar.

## Data availability

Raw sequencing has been deposited in the Gene Expression Omnibus (GEO) under accession number:xxxx. The global gene expression information of *B. thetaiotaomicron* during the time-course of colonization in both Transwell compartments (Figs. 3, 4) as well as in the *in*-*vitro* iron-depletion experiment (Fig. 5) was deposited on our public database, Theta-Base ([28, 29] accessible at http://bacteroides.helmholtz-hzi.de).

## ACKNOWLEDGEMENTS

We thank Sarah Reichardt and Lisa Pfeuffer for excellent technical support and Eric Martens (University of Michigan Medical School) for bacterial strains. We are grateful to Martin Fraunholz, Wilma Ziebuhr, and Chase Beisel for helpful discussions. This work was funded by the European Research Council (ERC Starting Grant #101040214 to A.J.W.), the German Research Foundation (DFG) through an Individual Research Grant (We6689/1-1 to A.J.W.), the SFB1583 DECIDE (#492620490; project A04 to A.J.W.), NIH NIAID (#F31AI178950, to R. T. F), NIH NIGMS (1R35GM147470, to W. Z.), NIH NIDDK (1R01DK134692, to W. Z.), Colorectal Cancer Alliance (#10065978, to W. Z.), Pew Charitable Trust (#2023-A-26048, to W. Z.), and The G. Harold & Leila Y. Mathers Charitable Foundation (MF-2207-03128, to W. Z.).

